# Geographic structuring of genetic variation differs across two contact zones in the *Diglossa carbonaria* superspecies

**DOI:** 10.64898/2026.07.14.738252

**Authors:** Anna E. Hiller, Brant C. Faircloth, Robb T. Brumfield

**Affiliations:** Department of Biological Sciences and Museum of Natural Science, Louisiana State University, Baton Rouge, LA 70803, USA

**Keywords:** RADseq, hybrid zones, gene flow, Andes, Thraupidae

## Abstract

Rapid radiations, where species quickly diversify with little to no change in the genetic composition of the taxa, provide unique opportunities to understand the processes underlying lineage divergence, especially when the newly formed species come into contact. This study examines three parapatrically distributed taxa of the Andean *Diglossa carbonaria* superspecies that differ strikingly in plumage and have been classified as an example of rapid lineage divergence. Using RADSeq data, we characterized the geographic structuring of genetic variation and investigated the possibility of introgressive hybridization at the contact zones between *D. humeralis atterima* and *D. b. brunneiventris* in northern Peru and between *D. b. brunneiventris* and *D. carbonaria* in Bolivia. We found weak genetic differentiation between *D. humeralis atterima* and *D. b. brunneiventris* and low, but diagnosable, differentiation between *D. b. brunneiventris* and *D. carbonaria*. At the contact zone between *D. humeralis atterima* and *D. b. brunneiventris*, the lack of genetic differentiation did not allow us to determine if hybridization was occurring between the taxa. In contrast, clustering analyses, diagnostic allele identification, and PCA analyses showed geographic patterns consistent with introgressive hybridization at the contact zone between *D. b. brunneiventris* and *D. carbonaria*, a geographic region where birds that are intermediate in plumage have been observed. Finally, we found a genetic break within the distribution of *D. b. brunneiventris*. The genetic divergence across this ∼450 km wide break is greater than that found between *D. humeralis atterima* and *D. b. brunneiventris*. Our results add to previous work documenting weak genetic differentiation between taxa that have striking plumage differences, and in showing that plumage color may be a poor phylogenetic marker.

**Lay Summary:**

- We studied three closely related avian taxa in Peru and Bolivia which have different plumages and adjacent ranges to examine how genetically distinct they are.
- To do this, we used RADseq data from 72 flowerpiercers sampled across their ranges to examine their population structure and to test whether the three taxa hybridize where their ranges meet.
- Despite their striking differences in plumage color, the three species showed few genetic differences. In one area where the ranges are adjacent, the groups were so similar that interbreeding could not be confirmed. In another, there was evidence of admixture, supported by observations of birds with intermediate plumage.
- For these taxa, feather color may not reflect evolutionary relationships, and more extensive genetic sampling is needed to understand how these plumage differences evolved.

## INTRODUCTION

Among montane ecosystems, the Andes contain some of the most species-rich communities in the world (Myers et al. 2000, Kier et al. 2009). This elevated diversity is particularly true for birds, where the Andes contain roughly 1/5th of avian species (Fjeldså and Krabbe 1990). Much of this Andean diversity is driven by the erosion and subsequent fragmentation of the mountains that isolate populations (Antonelli et al. 2018). Deep river valleys, for example, can act as dispersal barriers that lead to the geographic isolation of populations inhabiting ridges on either side of them (Remsen 1984, Graves 1985, 1988).

One phenotypically diverse avian group inhabiting high-elevation montane forests and grasslands of Mexico, Central, and South America is the flowerpiercers in the genus *Diglossa*. Flowerpiercers are a charismatic clade of 18 tanager species best known for their narrow, hooked bills, which they use to rob nectar from flowers (Vuilleumier 1969, Schondube and Martinez del Rio 2003). Within the flowerpiercer clade, the *Diglossa carbonaria* superspecies consists of four parapatrically and allopatrically distributed species (*D. carbonaria, D. brunneiventris, D. humeralis, and D. gloriosa*) found in the Andes of South America (Howard et al. 1991; Figure 1). Taxonomic studies have treated this group as four different species (Graves 1982), a single biological species (Zimmer 1929, Hellmayr 1935, Meyer de Schauensee 1970, Isler and Isler 1987), or a superspecies containing two semispecies (*D. humeralis* containing *D. h. humeralis, D. h. aterrima,* and *D. h. nocticolor*; *D. carbonaria* containing *D. c. carbonaria, D. c. brunneiventris*, and *D. c. gloriosa;* Vuilleumier 1969). For the purposes of this study, we treat the *D. carbonaria* superspecies as containing four biological species (*D. carbonaria, D. brunneiventris, D. humeralis,* and *D. gloriosa*) and seven subspecies (Figure 1).

**Figure 1.**
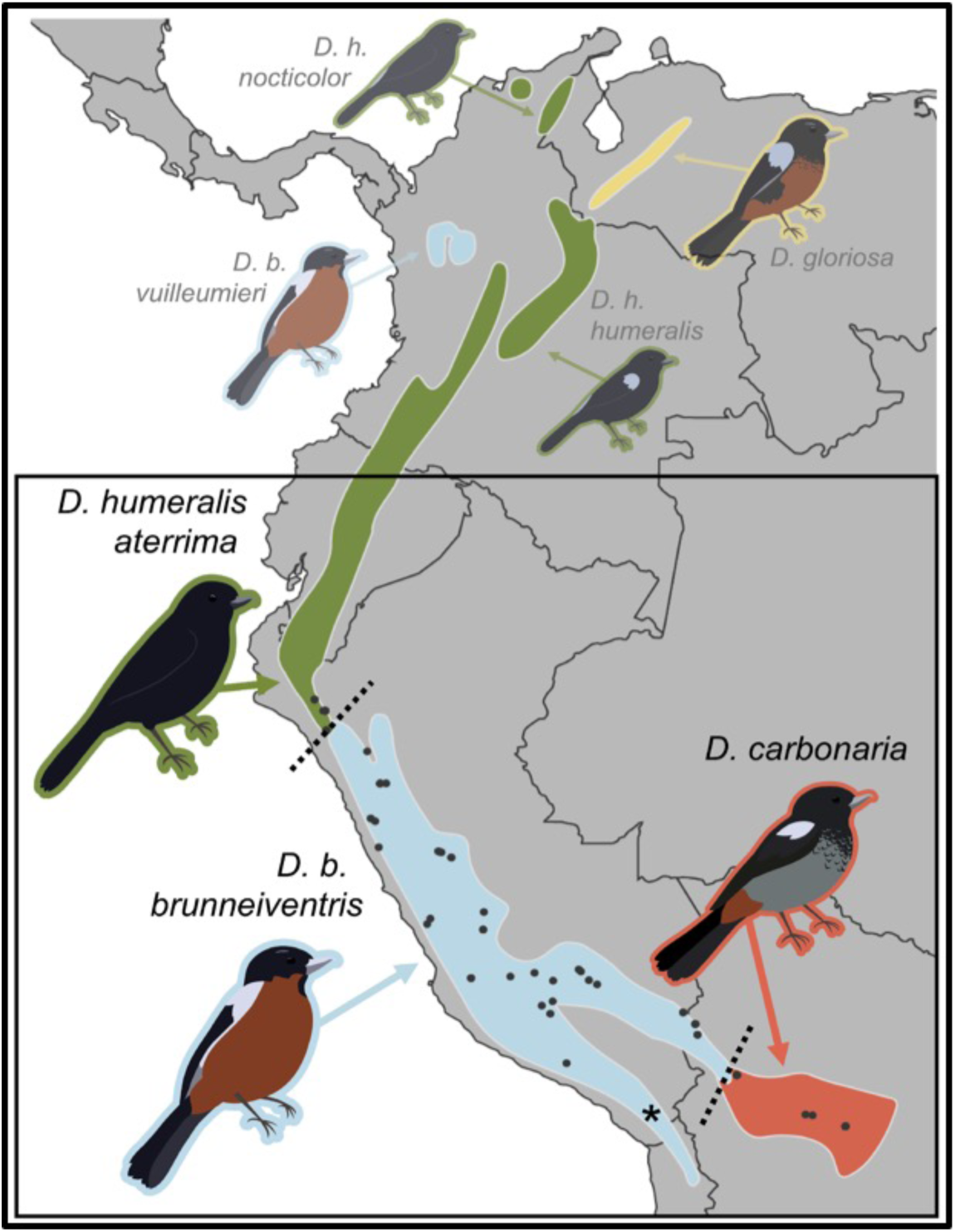
The *Diglossa carbonaria* superspecies, which includes four species and seven subspecies. Range maps are colored by species, and this color scheme carries throughout the remaining figures. Sampling points of individuals included in this study are indicated with black dots, and the two contact zones where there is a phenotypic transition between taxa are designated by dotted lines. The sampling point indicated with an asterisk (*) designates the collection locality of two *D. b. brunneiventris* individuals on the western slope of the Andes that our PCA results suggested were genetically distinct from the rest of the population.

The *D. carbonaria* superspecies is remarkable among flowerpiercers because it exhibits low genetic divergence but high divergence in plumage color and patterning among taxa (Figure 1). Mauck and Burns (2009) estimated that plumage divergence within the superspecies occurred during the last 500K years. A follow-up study (Burns et al. 2014) confirmed that taxa in the *D. carbonaria* superspecies diverged rapidly and recently, resulting in dramatic plumage differentiation yet little differentiation in mitochondrial DNA, similar to radiations of other tanagers, such as *Geospiza* finches (Lamichhaney et al. 2015) and *Sporophila* seedeaters (Turbek et al. 2021).

Rapid differentiation can make plumage an unreliable marker for indicating phylogenetic relationships (Brumfield et al. 2001), and convergence can occur if plumage differentiation is occurring in the context of evolutionary and physiological constraints that limit the palette of possible plumage character states (Stoddard and Prum 2011). Using complete taxon sampling of the *D. carbonaria* superspecies and multiple individuals per subspecies, Gutiérrez-Zuluaga et al. (Gutiérrez-Zuluaga et al. 2021) found a disconnect between plumage, species limits, and phylogenetic relationships. With the caveat that the data were limited to two mitochondrial genes (CYTB and ND2) and two nuclear introns, the authors showed that *D. brunneiventris* and *D. humeralis* were polyphyletic, providing additional evidence that plumage is a poor indicator of phylogenetic relationships (Gutiérrez-Zuluaga et al. 2021). For example, they discovered that *D. brunneiventris vuilleumieri*, a dark-backed, chestnut-bellied taxon inhabiting the Western and Central Andes of Colombia (Figure 1), was sister to *D. humeralis humeralis*, an all-dark taxon from the Eastern Andes of Colombia. Additionally, they found the all-dark taxon *D. humeralis aterrima* from the Andes of southern Colombia, Ecuador, and northern Peru was sister to *D. b. brunneiventris*, a dark-backed, chestnut-bellied taxon that replaces *D. h. aterrima* in northern Peru and which extends south to northern Bolivia, where it is replaced by *D. carbonaria*, which has a dark back, gray belly, and chestnut crissum (Figure 1).

Within the *D. carbonaria* superspecies, the three taxa *D. brunneiventris vuilleumieri* (dark-backed, chestnut-bellied), *D. humeralis aterrima* (all dark), and *D. brunneiventris brunneiventris* (dark-backed, chestnut-bellied) exhibit a “leapfrog” pattern of geographic variation (Figure 1), which Remsen (1984) defined as geographic variation in which two disjunct taxa are more similar in plumage to each other than to a third geographically intermediate taxon in the same species complex. In addition to the *D. carbonaria* superspecies, Remsen (1984) identified leapfrog variation in the distantly related *Diglossa lafresnayii* superspecies, which exhibits a remarkably similar pattern to the *D. carbonaria* superspecies: two disjunct, all-dark taxa (*D. lafresnayii* and *D. mystacalis*) occur on either side of a dark-backed, chestnut-bellied taxon (*D. gloriosissima*). Mauck and Burns (2009) suggested that these two instances of leapfrog variation in *Diglossa* (i.e., in the *D. carbonaria* and *D. lafresnayii* superspecies) likely originated at different points in time. Collectively, prior phylogenetic analyses and observations of plumage differences suggest that plumage evolution occurs rapidly in *Diglossa* and that a limited palette of plumage colors and plumage patterns may produce these superficially similar, ‘convergent’ plumages among distantly related *Diglossa* taxa.

Where different taxa of *Diglossa* come into contact, such as between *D. humeralis aterrima* (all dark) and *D. brunneiventris brunneiventris* (dark-backed, chestnut-bellied) in northern Peru, the contact has been assumed to be secondary (Graves 1982). An alternative hypothesis is that the contact zones represent the termini of recently derived and actively spreading plumage traits. For example, if an ‘all-black’ plumage allele arose and spread within the northern distribution of *D. b. brunneiventris*, its contact zone with *D. humeralis aterrima* might represent the southern terminus of that spreading allele. Conversely, if the ‘dark-backed, chestnut-bellied’ allele arose somewhere within the distribution of *D. b. brunneiventris* or *D. carbonaria*, the *D. humeralis aterrima* x *D. b. brunneiventris* contact zone represents the northern terminus of that allele (Figure 1). Gutiérrez-Zuluaga et al. (2021) found *D. humeralis aterrima* and *D. b. brunneiventris* to be genetically indistinguishable, so either of these explanations is possible. And, these types of plumage evolution events across contact zones have been reported in other avian genera. For example, the yellow throat of the Golden-collared Manakin (*Manacus vitellinus*) arose relatively recently from a white-plumaged ancestor and is actively spreading from its origin in Panama into populations in northwestern Colombia and eastern Costa Rica (Parsons et al. 1993, Lim et al. 2024). At the vanguard of the spreading yellow trait in Colombia, where yellow-throated and white-throated birds come into contact, individuals appear to be genetically indistinguishable except for the allele(s) underlying the yellow trait (Brumfield et al. 2001, Brumfield and Carling 2010, Harvey et al. 2020, Lim et al. 2024).

Genetic analysis of contact zones in groups like the *D. carbonaria* superspecies can provide insights into the evolutionary and ecological mechanisms associated with the origin and maintenance of genetic and phenotypic characters (Harrison 1993). The first study (Zimmer 1929) of the contact zone between *D. humeralis atterima* and *D. b. brunneiventris* in northern Peru (Figure 1) noted the absence of intermediate-plumaged individuals. Using a larger number of specimens, Vuilleumier (1969) concluded that *D. humeralis atterima* and *D. b. brunneiventris* were completely reproductively isolated. Graves (1982) sampled the first transect across the same contact zone and found no evidence of ‘hybrid-like’ individuals. This observation was especially notable because he observed mixed pairings between female *D. b. brunneiventris* and male *D. humeralis atterima* during his fieldwork. Graves (1982) also noted that aspects of the habitat and landscape in the region where the distributions of *D*. *humeralis aterrima* and *D. b. brunneiventris* meet greatly diminish contact between the two taxa. For example, the corridor of appropriate shrubland habitat through which birds could disperse was only about 3 km in width where they came into contact (Graves 1982). Given the lack of observations of nests or fledglings from the territories of the mixed pairings and the limited opportunities for contact imposed by the landscape, Graves (1982) concluded that while there may be occasional interbreeding between *D. humeralis atterima* and *D. b. brunneiventris*, the two taxa should be treated as distinct biological species.

At the Bolivian contact zone between *D. b. brunneiventris* and *D. carbonaria* (Figure 1), Zimmer (1929) described several individuals with intermediate plumage. Carriker (1935) noted that the two taxa could be found together at a locality ‘a few miles’ south of La Paz and that there was no evidence of intergradation. Niethammer (1956) subsequently noted several specimens collected 40 km ESE of La Paz that showed clear intermediacy in plumage, and Vuilleumier (1969) used these records and additional specimen collections to conclude that the two taxa were experiencing some, though limited, introgressive hybridization. Graves (1982) re-examined the Vuilleumier specimens and concurred that “a narrow zone of hybridization between *D. b. brunneiventris* and *D. carbonaria* occurs northeast of La Paz” as well as that the location of the hybrid zone was largely concordant with the ecological transition from wetter to drier montane scrub forest. Graves made special note of an unusual pattern in which hybrid-like *D. b. brunneiventris* plumage characters were found throughout the distribution of *D. carbonaria* and, surprisingly, were highest in frequency in populations of *D. carbonaria* farthest from the hybrid zone. Graves (1982) suggested that this plumage polymorphism did not reflect introgression from *D. b. brunneiventris* into *D. carbonaria*, and he proposed an alternative hypothesis that retained ancestral plumage alleles explained these observations. However, these observations were based on a relatively small sample size especially from the putative contact zone.

Here, we use RADseq data collected from vouchered tissue samples of *D. humeralis atterima, D. b. brunneiventris,* and *D. carbonaria* to characterize the geographic structure of genetic variation in these populations. We test for evidence of introgressive hybridization across the Peruvian contact zone between *D. humeralis atterima* and *D. b. brunneiventris* and the Bolivian contact zone between *D. b. brunneiventris* and *D. carbonaria*. At each contact zone, we assess whether hybridization is occurring, we examine the geographic extent of introgression, and we test the hypothesis that the geographic location of plumage transitions between subspecies coincide with the locations of the genetic transitions between subspecies. Discrepancies between the genetic and plumage transition locations could support a hypothesis of unidirectional introgression of plumage traits or of a spreading plumage allele. We also test for the presence of a genetic break within the distribution of *D. b. brunneiventris*, as recently reported (Gutiérrez-Zuluaga et al. 2021).

## METHODS

### Sampling

We sampled 72 individuals of the *D. carbonaria* superspecies from northern Peru to central Bolivia, including samples near the contact zone between *D. humeralis atterima* and *D. b. brunneiventris* in northern Peru and between *D. b. brunneiventris* and *D. carbonaria* in northern Bolivia (Figure 1). By straight line distance, our southernmost sample of *D. humeralis atterima* and our northernmost sample of *D. b. brunneiventris* were collected approximately 150 km apart, and our southernmost sample of *D. b. brunneiventris* was collected 215 km from our northernmost sample of pure *D. carbonaria,* with a few individuals with *D. carbonaria*-like plumage collected from the region where the two populations come into contact (∼50km south of the nearest *D. b. brunneiventris* sample). Because we were primarily interested in assessing the possibility of introgressive hybridization across these two contact zones, we did not include samples of other subspecies of *D. humeralis* or *D. brunneiventris* from the northern part of the superspecies’ range in Ecuador, Colombia, or Venezuela (Figure 1). We also included two outgroup individuals of *D. albilatera,* which represent the sister group to the *D. carbonaria* superspecies, to root the trees produced by SNAPP analyses. The total of 74 tissues included seven samples we collected in the field and 64 samples we received as loans from other institutions (Table S1). At the time of collection, tissues were flash-frozen in liquid nitrogen or preserved in 95% ethanol.

### Restriction site-associated DNA sequencing (RADseq)

We extracted total DNA from each tissue using the Qiagen DNeasy Blood and Tissue Kit following the manufacturer’s instructions. We quantified DNA extracts using a Qubit fluorometer (Life Technologies, Inc.), and we standardized concentrations of working samples to 10ng/µl. During subsequent lab steps, we included one “blank” (water) sample as a negative control for cross-contamination checks. After preparing extracts, we collected RADseq data from the 74 individuals following the 3RAD protocol (Bayona-Vásquez et al. 2019) and using the enzymes NheI-HF, EcoRI, and XbaI. After RADseq library construction, we pooled all samples, attached a pair of iTru sequence tags to the pool (Glenn et al. 2019) using 12 PCR cycles, and used a column cleanup to remove small DNA fragments. We combined the 74 RADseq libraries with 22 additional RADseq libraries that were not part of this study (having different sequence tags) and sequenced the 96 samples across two lanes of paired-end (PE) 150 base-pair sequence on an Illumina NovaSeq 6000 (University of Kansas Medical Center Genomics Core) targeting a total of 200 M reads for the pool of RADseq libraries from this study.

### Single nucleotide polymorphism (SNP) calling and filtering

We demultiplexed, trimmed, and cleaned raw reads in Stacks v2.65 (Catchen et al. 2011, 2013, Rochette et al. 2019) with the “*process_radtag*” command. Then, we aligned these data to a reference genome from a female *D. b. brunneiventris* collected in central Peru (GCA_019023105.1, Hiller et al. 2021) using BWA v0.7.17-r1188 (Li and Durbin 2009, Li 2013). We used samtools v1.1 (Danecek et al. 2021) to remove imperfect matches, reads with mapping quality <25, reads with >5 SNPs, and to sort the resulting BAM files. Then, we used “*gstacks*” to build RAD loci from reads aligned to the reference genome and to call SNPs. We converted the resulting data into a variant call format (VCF) file using the “*populations*” command, and we specified samples as belonging to four populations in a “*population map*” file based on the taxonomic identification of each specimen in its corresponding collection (*D. humeralis atterima*, *D. b. brunneiventris*, *D. carbonaria*, and *D. albilatera,* Table S1). We ran Stacks using default parameters unless otherwise specified.

Because the quality of SNP calls can greatly affect the downstream inference of population genetic structure (Chattopadhyay et al. 2014, Linck and Battey 2019), we used vcftools v1.17 to filter the VCF file by removing loci having a minor allele count less than three (*--mac* 3), loci with less than 15x depth of coverage (*--min-meanDP* 15), as well as indels and sites that were not bi-allelic (*--remove-indels --min-alleles* 2 *--max-alleles* 2), and loci that were missing SNP calls from more than 25% of individuals (*--max-missing* 0.75).

Then, we removed loci that were out of Hardy-Weinberg equilibrium (HWE, Wigginton et al. 2005) by dividing the VCF file by the taxonomic identification of each species, removing loci out of HWE for each taxonomic group (*--hwe* 0.05), and recombining the filtered groups into a single VCF file. After recombining, we again filtered the resulting file to ensure loci were biallelic and that depth of coverage, minor allele count, and data missingness remained the same. We ran additional steps to remove sites that were potentially in linkage disequilibrium (*--thin* 10000) as well as those sites having low mappability, which we computed using a genmap v1.3.0 (Pockrandt et al. 2020) analysis of the reference genome sequence. Finally, we used the *--missing-indv* function of vcftools to ensure that all individuals retained in the data set had genotype calls for ≥75% of sites. We refer to this file as the primary VCF file.

### Species tree

To visualize the degree and position(s) of potential uncertainty in the relationships among our samples, we used the coalescent-based species tree inference program SNAPP (Bryant et al. 2012) in Beast 2 (Bouckaert et al. 2019). Because our dataset was large and SNAPP is computationally intensive, we created two subsets of data that included one outgroup individual (*D. albilatera*) as well as ten ingroup individuals: two from each taxon at the periphery of the range (*D. humeralis atterima* and *D. carbonaria*) and four from the intervening taxon (*D. b. brunneiventris*), preferentially including individuals that had fewer missing genotypes. For both sets, we inferred the posterior distribution of species trees with a Markov chain Monte Carlo (MCMC) chain length of 5 million, we sampled trees every 1,000 iterations, and we discarded 1% (50,000 iterations or 50 sampled trees) as burnin. To assess convergence, we visualized outputs in Tracer v1.7.1 (Rambaut et al. 2018) and ensured ESS values for most parameters were >200. We then visualized the posterior distribution of trees for each run using DensiTree (Bouckaert 2010) after rooting trees on the outgroup (*D. albilatera*).

### Population structure

To characterize the geographic structuring of genetic variation, we calculated ancestry coefficients for individuals using PopCluster v1.5.0.0 (Wang 2022, 2024), running separate analyses with K values of 1 to 3, and selecting the “best” K using the D_LK2_ estimator. We also visualized genetic structure in the dataset using a principal components analysis (PCA) implemented in SNPRelate (Zheng et al. 2012). We estimated the distance of samples from their respective contact zones by excluding individuals collected in contact zones and defining the contact zone center in Peru for *D. humeralis atterima x D. b. brunneiventris* at the Cachula ravine (-6.9, -78.52), approximately halfway between our southernmost sample of *D. humeralis atterima* and our northernmost sample of *D. b. brunneiventris*. Likewise, for *D. b. brunneiventris x D. carbonaria*, we defined the contact zone center as the town of Tres Rios, Bolivia (-16.57, -67.79), halfway between our southernmost sample of *D. b. brunneiventris* and northernmost sample of *D. carbonaria*. This ‘center’ designation is for convenience and should not be interpreted beyond that, because our sampling is insufficient to discern the finer-scale structure of the geographic transition from one taxon to the other.

### Geographic patterns of diagnostic alleles

The population structure analyses suggested that genetic breaks exist between groups of individuals, but none of these breaks corresponded with the contact zone between *D. humeralis atterima x D. b. brunneiventris* in Peru. As a result, we were interested in examining transitions in individual SNPs, so we identified “diagnostic alleles” and examined their geographic structuring. Specifically, we used *adegenet* (Jombart 2008) to calculate allele frequencies of each SNP, and we considered SNPs with allele frequencies ≤0.2 in *D. b. brunneiventris* and ≥0.8 in *D. humeralis atterima* as diagnostic for the contact zone in Peru while we considered SNPs with allele frequencies ≤0.2 in *D. b. brunneiventris* and ≥ 0.8 in *D. carbonaria* as diagnostic for the contact zone in Bolivia. We were interested in whether the frequency of diagnostic SNPs gradually decayed as a function of geographic distance from the contact zone, which would be a pattern more consistent with introgressive hybridization, or instead showed a more random geographically unstructured distribution of shared alleles consistent with incomplete lineage sorting. To examine this pattern, we extracted the genotypes for each of the diagnostic alleles in each respective group from the VCF file and plotted these in geographic space. We refer to these diagnostic SNPs using the first component of the ID column in the primary VCF file, which is a unique integer value assigned to each locus by stacks.

## RESULTS

### Sequencing and SNP quality

We collected an average of 1,029,412 (SD 623,168) PE150 sequence reads per individual, and our Stacks analysis identified 123,585 variable loci having a mean, per-sample coverage of 30x (standard deviation 17x, range 6-85x). Stacks identified 411,876 variant sites across these loci, and filters to remove variants that were not biallelic, had coverages <15x, or that were missing variant calls in ≥25% of individuals reduced this number to 49,985. Additional filters to remove variants that were not in HWE by taxonomic group, in poorly mapping regions of the genome, or potentially in linkage disequilibrium reduced this number to 6,675. After applying these filters, five individuals fell below our threshold for completeness (75% of sites present in all individuals), and we removed them. This produced a primary data set that included 6,675 biallelic site for 69 individuals (67 ingroup individuals and 2 outgroup individuals, Table S1).

### SNAPP analysis

After creating two subsets of individuals, the SNAPP analyses included 3286 SNPs (run1) and 3102 SNPs (run2). The lower numbers of SNPs relative to the primary VCF file resulted from the fact that SNAPP removed sites having missing genotypes. The analyses appeared to converge after 5 million generations based on visualization of the log files in Tracer, with ESS values for all parameters (posterior, likelihood, prior, u, v) >200 except in one case (run1 ESS for u was 188). Both SNAPP runs produced similar topologies (Figure 2, Figure S1), suggesting the individuals representing *D. humeralis atterima* were nested within *D. b. brunneiventris* and calling into question the monophyly of both taxa. The cloudogram of posterior trees also showed areas of conflicting signal that were associated with *D. b. brunneiventris* individuals collected near the contact zones in Peru and Bolivia, a pattern consistent with introgression or incomplete lineage sorting.

**Figure 2.**
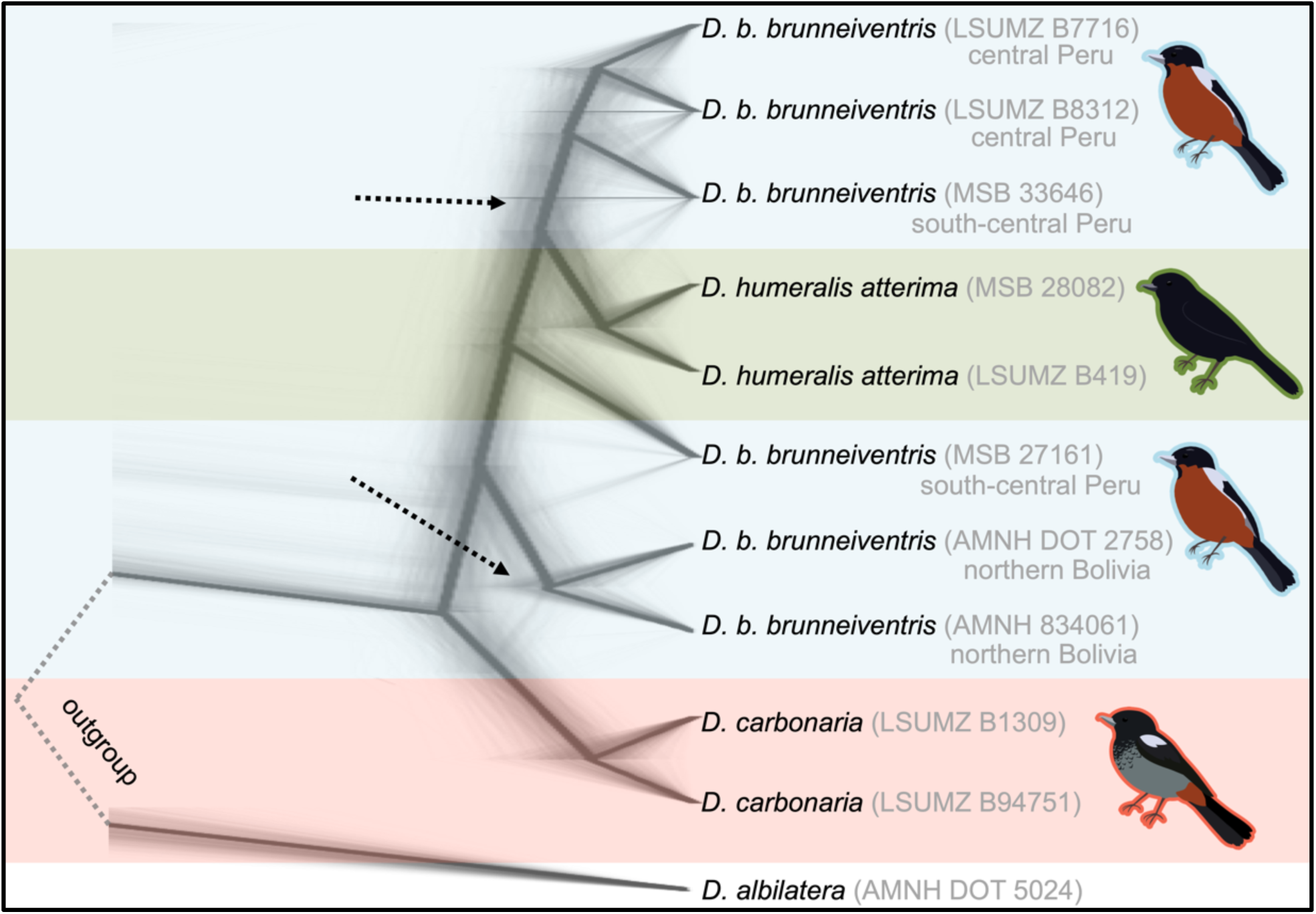
Densitree plots of one SNAPP analysis illustrating the relationships of individuals from each sampled population in the *D. carbonaria* superspecies, with *D. albilatera* as an outgroup (Supplemental Figure S1 illustrates results from the second SNAPP analysis). The results show the cloudogram of posterior trees, with the density (or darkness) of the cloudogram indicating how often a particular topology occured. Alternate topologies suggest uncertainty in the tree, particularly where indicated by the dotted arrows, which is a pattern consistent with introgression or incomplete lineage sorting.

### Geographic structure of genetic variation

Based on the DLK2 estimator, the optimal number of genetic clusters in the dataset was K=2, with a genetic break occurring at the contact zone in Bolivia on the eastern slope of the Andes near La Paz, separating *D. carbonaria* individuals from all other individuals (*D. b. brunneiventris* and *D. humeralis atterima*) we sampled (Figure 3). When we increased K to 3, matching current taxonomy, this break remained. Admixture proportions across both analyses were largely similar. At a K=3 four of six (67%) individuals with the *D. b. brunneiventris* phenotype collected within 280 km NNW of the contact zone center showed a moderate proportion of *D. carbonaria* ancestry (0.195 to 0.26), while one of six (16%) individuals with the *D. carbonaria* phenotype collected within the contact zone showed a low proportion (0.128) of *D. b. brunneiventris* ancestry.

**Figure 3.**
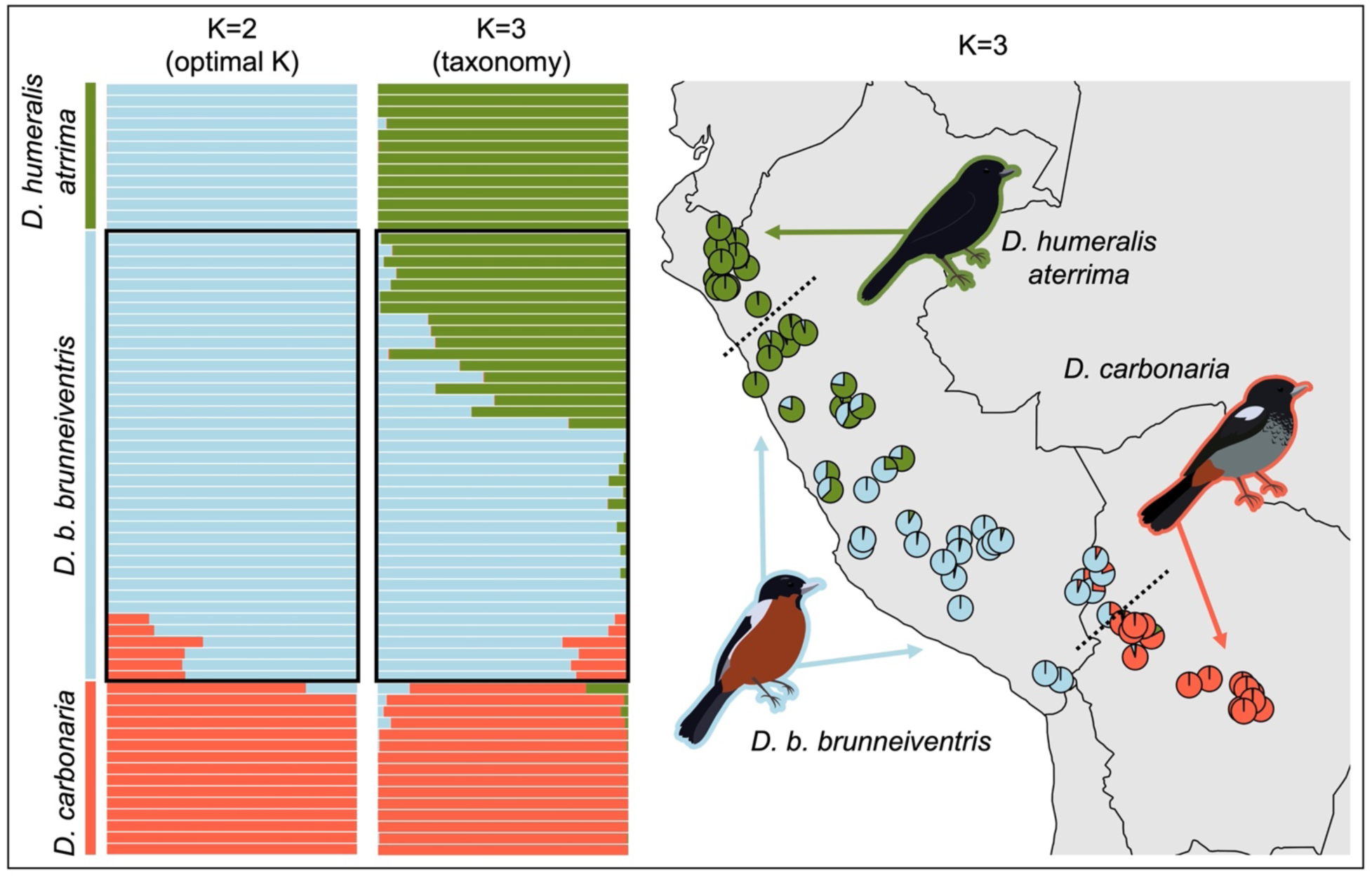
With a K of 2 (the optimal value based on the D_LK2_ estimator, left-most bar chart) there is a genetic break corresponding to the contact zone between *D. b. brunneiventris* and *D. carbonaria* (lower dotted black line in Bolivia). With a K of 3 (the number of clusters matching taxonomy and major phenotypic differences, right-most bar chart), we detected an additional genetic break in central Peru within the distribution of *D. b. brunneiventris*. Note that this location is well outside of the expected transition zone between plumage phenotypes (illustrated by the upper dotted black line in northern Peru). The black boxes surrounding the bar charts indicate individuals identified as *D. b. brunneiventris* in collections based on plumage phenotype. All individuals are organized north-to-south in both the bar charts and the map. For the map, ancestry proportions for each individual (at K=3 to match taxonomy) are represented as pie charts and plotted on a map of western South America by locality (jittered to minimize visual overlap).

We also detected a second genetic break when we increased K to 3 that corresponded to a geographic position in the middle of the range of *D. b. brunneiventris –* a location 700 km away from the contact zone in Peru where the phenotypic transition between *D. b. brunneiventris* and *D. h. aterrima* occurs. Although birds on either side of this genetic break are phenotypically indistinguishable, the break suggests that individuals with *D. b. brunneiventris* phenotypes north of this location are genetically more similar to *D. humeralis atterima* than they are to *D. b. brunneiventris* south of this location. Individuals sampled from ∼185 km northwest to ∼245 km southeast of the contact zone exhibited similar admixture proportions (0.924 - 0.963 *D. humeralis atterima* ancestry) despite those individuals collected north of the contact zone having different phenotypes from those collected south of the contact zone.

The PCA (Figure 4) largely corroborated the PopCluster analysis (Figure 3). The first principal component explained 4.57% of the variation and separated *D. humeralis atterima* and *D. b. brunneiventris* individuals from *D. carbonaria* individuals. Variation along PC1 was more extensive among *D. carbonaria* individuals than among individuals of *D. humeralis atterima* and *D. b. brunneiventris*. The second principal component, which explained 2.25% of the variation, captured differences in the *D. humeralis atterima* and *D. b. brunneiventris* individuals. Specifically, *D. humeralis atterima* individuals clustered together on one end of the PC2 axis but transitioned into *D. b. brunneiventris* individuals – a pattern consistent with the slow transition that we observed in estimated admixture proportions between these two taxa. The PCA also suggested that two individuals with *D. b. brunneiventris* phenotypes were genetically quite different from most other samples representing this taxon (marked by an asterisk in Figure 4). These two samples were collected from the southernmost extent of the *D. b. brunneiventris* range, which extends in a narrow band along the western slope of the Andes into northern Chile, physically separated from contact with *D. carbonaria* by the Altiplano plateau.

**Figure 4.**
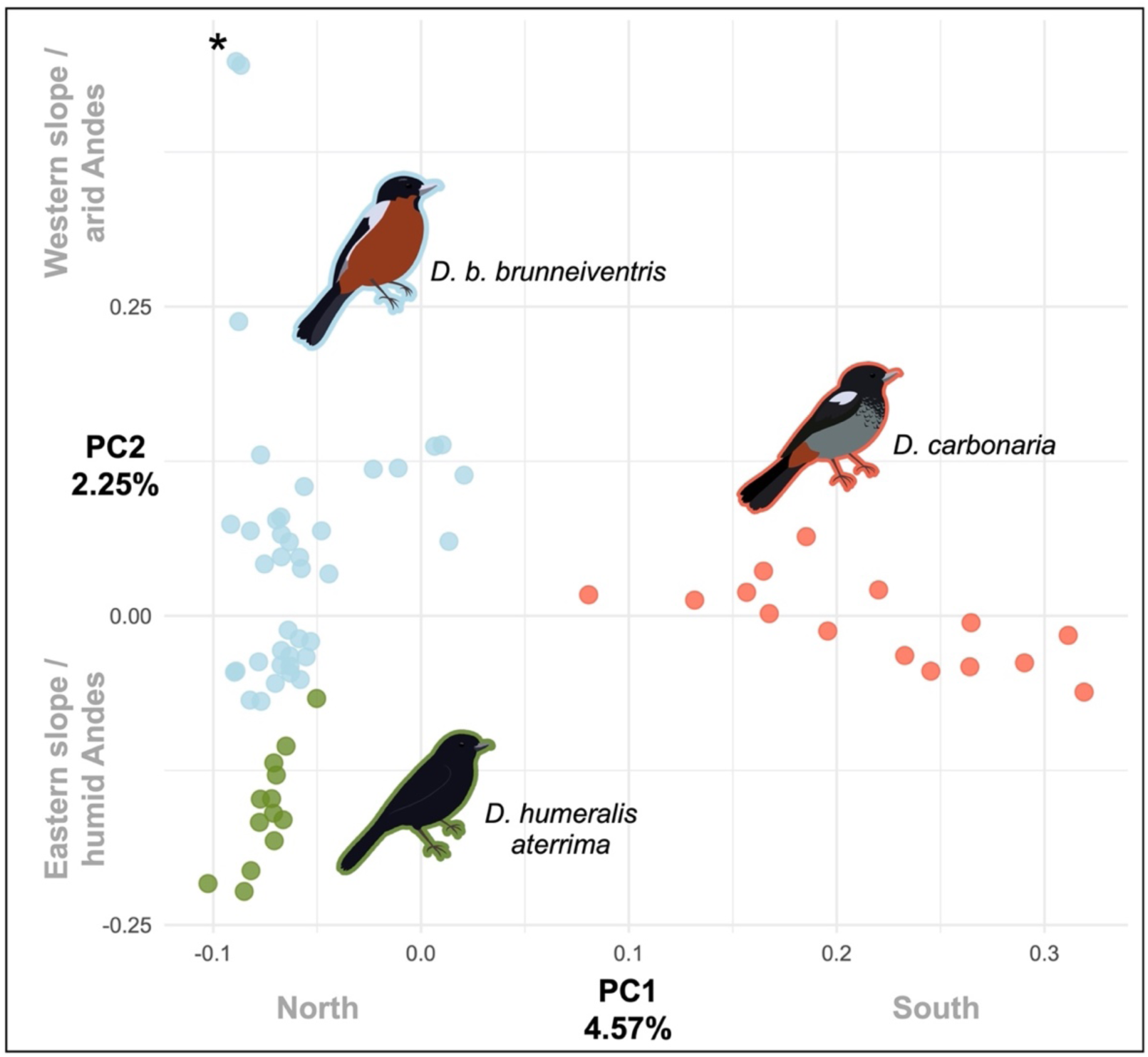
Principal components analysis of genetic differences among the samples collected with points colored to reflect their subspecies identification (the same color is used around the subspecies illustration). PC1 (x-axis) shows differentiation between taxa that generally corresponds to the position of their collection localities along a north to south axis while PC2 (y-axis) shows differentiation between the samples that generally corresponds to their position on an east (humid) to west (arid) axis across the Andes. The asterisk indicates the two *D. b. brunneiventris* individuals collected from the southernmost extent of their range, on the western slope of the Andes. Note that individuals of *D. humeralis atterima* cluster together on PC2 but grade into individuals of *D. b. brunneiventris*.

### Diagnostic alleles

Allele frequency analysis identified one diagnostic SNP in comparisons of *D. b. brunneiventris* to *D. humeralis atterima*. Geographic mapping of genotypes at this locus (SNP 84746, Figure 4) showed that frequencies for the alternate allele were 0.86 in *D. humeralis atterima*, occurred at lower frequencies in *D. b. brunneiventris* collected from the northern part of its distribution, and occurred at very low frequencies in *D. b. brunneiventris* collected from the southern part of its distribution. This geographic pattern is generally inconclusive because it could be due to introgression of the alternate allele across a putative *D. humeralis atterima* x *D. b. brunneiventris* hybrid zone, but it also could reflect an ancestral polymorphism that was fixed due to drift or selection in *D. humeralis atterima*, or it could be explained by an isolation-by-distance effect if there is stepping-stone migration and no reproductive barriers exist between *D. humeralis atterima* and *D. b. brunneiventris*.

A similar analysis identified four diagnostic SNPs (12979, 88589, 99348, 107922) in comparisons of *D. b. brunneiventris* to *D. carbonaria* (Figure 4). At these loci, frequencies for the reference allele were high (0.88-1.0) among individuals having *D. b. brunneiventris* phenotypes and transitioned sharply to the alternate allele (frequencies 0.8-0.9) across the Bolivian contact zone in individuals having the *D. carbonaria* phenotype. At SNP 12979, we identified heterozygotes only among individuals collected within the contact zone. For the remaining SNPs, we sometimes detected heterozygotes in *D. b. brunneiventris* north of the contact zone and *D. carbonaria* south of the contact zone. The relatively narrow region of limited introgression at some loci (e.g., SNP 12979) is consistent with a hybrid zone between *D. b. brunneiventris* and *D. carbonaria*.

**Figure 5.**
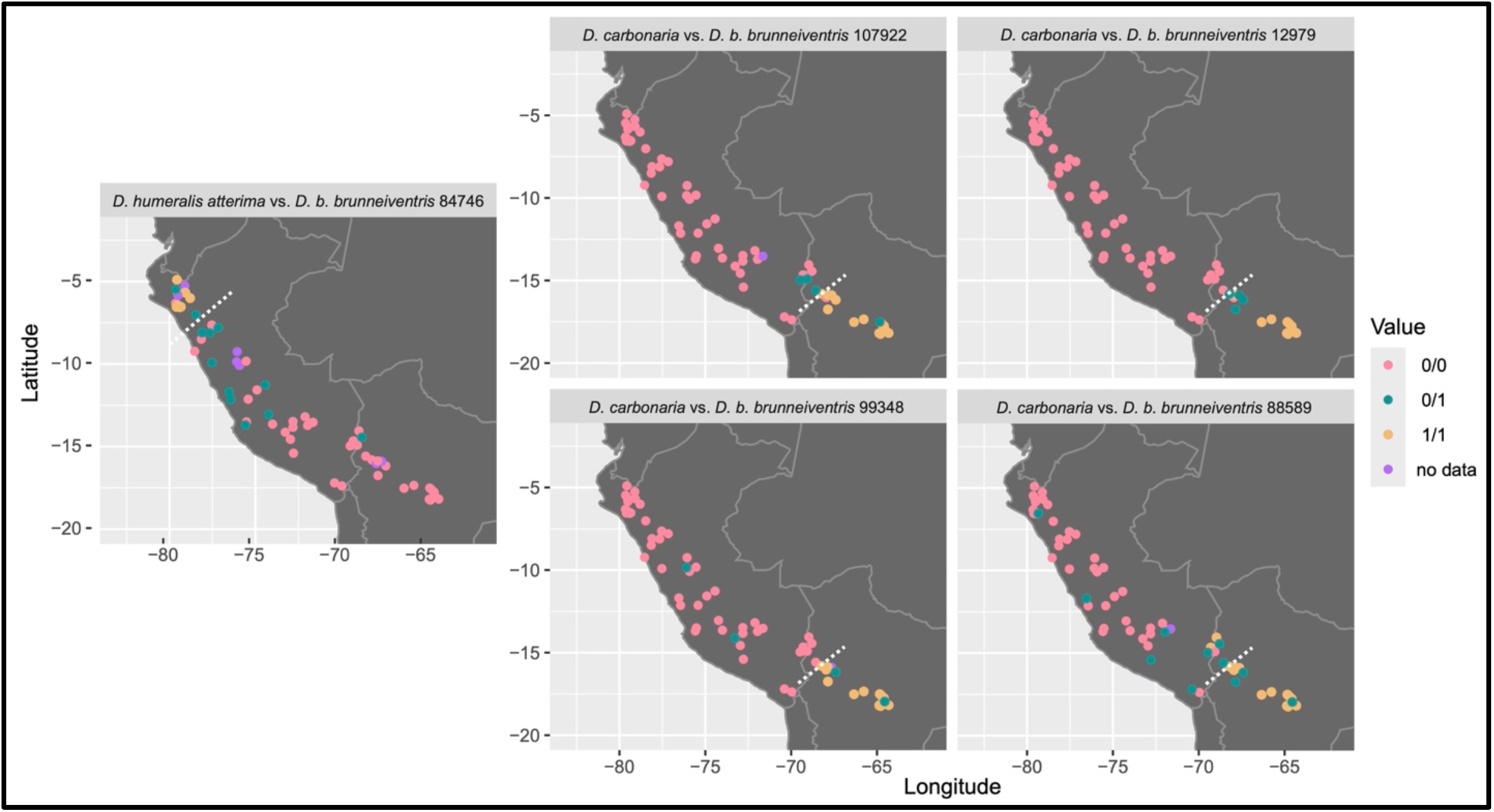
The spatial position of genotypes at loci that we identified as being “diagnostic” in our analyses (outside of an allele frequency range of 0.2-0.8) for the comparisons described in the panel headers. Four loci were diagnostic for *D. carbonaria* vs. *D. b. brunneiventris* and one locus was diagnostic for *D. humeralis atterima* vs. *D. b. brunneiventris*. Point colors correspond to the genotypes in the legend.

## DISCUSSION

Our data corroborate previous genetic studies of *Diglossa* (Mauck and Burns 2009, Gutiérrez-Zuluaga et al. 2021) by showing that the three taxa we examined from the *D. carbonaria* superspecies had low genetic differentiation (K=2, Figure 3) despite obvious differences in plumage color and patterning, providing additional evidence to the general idea that plumage color and pattern can be unreliable indicators of phylogenetic relationships (Brumfield et al. 2001). In the species trees we constructed (Figure 2, Supplemental Figure S1), *D. humeralis aterrima* (all dark plumage) was nested within a clade of *D. b. brunneiventris* and sister to individuals collected from the northern part of the *D. b. brunneiventris* (dark-backed, chestnut-bellied) range, calling into question the monophyly of both taxa. This same relationship was found by (Gutiérrez-Zuluaga et al. 2021), although our results suggest that the confusing placement of *D. humeralis* could be due to allele sharing with *D. b. brunneiventris* at the northern end of their range. Unfortunately, we cannot explain uncertainty in the tree because it could result from either introgression or incomplete lineage sorting. We also resolved the clade representing *D. carbonaria* as sister to all remaining individuals of *D. b. brunneiventris* and *D. humeralis aterrima* that we sampled. Again, the posterior sample of gene trees suggested the possibility of allele sharing between *D. carbonaria* and *D. b. brunneiventris* collected near the contact zone in the southern portion of their range that could be explained by introgression or incomplete lineage sorting.

### *Hybridization at the* D. humeralis aterrima *and* D. b. brunneiventris *contact zone?*

Our remaining analyses were equivocal with regards to hybridization occurring at the contact zone between *D. humeralis aterrima* and *D. b. brunneiventris*. PopCluster analyses at K=2 did not identify any genetic break between these phenotypically different taxa, and PopCluster analyses at K=3 did not identify a genetic break between these taxa associated with the contact zone. Similarly, the PCA revealed few genetic differences between individuals with *D. humeralis atterima* and *D. b. brunneiventris* phenotypes collected near the contact zone, and the amount of variation explained by the PC axis separating these taxa was low (2.25%). Relatively weak differentiation between *D. humeralis atterima* and *D. b. brunneiventris* in the PopCluster analyses and PCA could be due to their relatively recent divergence, to extensive introgression since secondary contact, or to both processes. The single diagnostic allele we detected between them was nearly fixed (frequency 0.86) in *D. humeralis aterrima*, but the allele also occurred at moderate frequencies throughout the northern distribution of *D. b. brunneiventris*. This geographic pattern could potentially indicate introgression, although it did not exhibit the expected decay in frequency moving southward from the contact zone (Figure 3). One potential issue in our study design that could confound our understanding of relationships between these two taxa is that we did not sample the complete range of *D. humeralis atterima*. From our northernmost sample, the distribution of *D. humeralis atterima* extends another 1000 km northward into Ecuador and Colombia. If there is a genetic break between *D. humeralis atterima* and *D. b. brunneiventris* that is north of our northernmost sample, we would have failed to detect it. For now, we conclude that despite the pronounced plumage differences between *D. humeralis atterima* (all black) and *D. b. brunneiventris* (dark-backed, chestnut-bellied), the amount of genetic differentiation between them is low and may be limited to loci associated with plumage differences.

If hybridization is occurring between *D. humeralis atterima* and *D. b. brunneiventris*, we hypothesize that the lack of obvious phenotypic hybrids between them may be an example of melanin-based color masking of the hybrid individuals, where one color covers another, often corresponding to black plumage (see Price-Waldman and Stoddard 2021 for a review). Examples of this phenomenon in birds are numerous (Nero 1954, Moreau 1958, M. Hofmann et al. 2007, Hudon et al. 2015, Aguillon et al. 2021). Under this hypothesis, plumage traits that might identify admixed *D. humeralis atterima* and *D. b. brunneiventris* are masked by melanin and hybrid individuals appear entirely black. Future work examining the degree of differentiation between the genomes of *D. b. brunneiventris* and *D. humeralis atterima* obtained from the zone of secondary contact could provide a good starting point to test this hypothesis.

### Hybridization at the D. b. brunneiventris and D. carbonaria contact zone

Consistent with previous reports (Vuilleumier 1969, Graves 1982) of phenotypically hybrid-like individuals in and around the contact zone between *D. b. brunneiventris* and *D. carbonaria,* our admixture and diagnostic allele analyses suggest that hybridization may be occurring between these two taxa. Admixture analyses at K=2 and K=3 identified a genetic break between these two taxa (Figure 3) with individuals of admixed ancestry occurring on either side of the contact zone. Admixture proportions suggest the presence of a hybrid zone east of La Paz that is relatively wide in terms of genetic introgression (∼300 km). The diagnostic alleles that we identified generally support the admixture results – the relatively narrow regions of limited introgression at these loci are consistent with a hybrid zone between *D. b. brunneiventris* and *D. carbonaria*. The amount of introgression from *D. b. brunneiventris* into *D. carbonaria* we found though is too limited to explain the geographic distribution of the plumage polymorphism within *D. carbonaria* described by Graves (1982) where *D. carbonaria* individuals with hybrid-like characters were found to the south of any area of potential contact with *D. b. brunneiventris*. Past introgressive hybridization or retained ancestral polymorphism remain viable alternative hypotheses for this observation.

### Genetic differences in Diglossa do not necessarily correspond to plumage differences

One of the hallmarks of the *Diglossa carbonaria* superspecies is that plumage differentiation can occur rapidly (Burns et al. 2014). However, we identified two cases of genetic differentiation that was not associated with plumage differentiation. First, we identified a ∼450 km-wide zone of genetic differentiation in central Peru within the distribution of *D. b. brunneiventris* (Figure 3) that generally corresponds with a phylogeographic break detected by Gutiérrez-Zuluaga et al. (2021) in their mitochondrial data (Figure 4). Could this break represent the true genetic contact zone between *D. humeralis aterrima* and *D. b. brunneiventris*, with the northern contact zone between different phenotypes reflective of introgressing plumage traits? Other hybrid zone studies have found instances of displaced genetic and phenotypic clines (e.g., Parsons et al. 1993), but to the best of our knowledge, a displacement of 700 km between genetic and phenotypic clines in *Diglossa* would be unprecedented in terms of distance. The second case we observed was between individuals of *D. b. brunneiventris* collected from the western versus the eastern slope of the Andes. Specifically, we found more separation between these individuals of *D. b. brunneiventris* on the second PCA axis (Figure 4) than we identified between individuals of *D. humeralis atterima* and *D. b. brunneiventris*, suggesting taxonomy in this group needs revision.

### The “leapfrog” pattern in the D. carbonaria superspecies

Our data may provide insights into the leapfrog pattern Remsen (1984) described in the *D. carbonaria* superspecies. The leapfrog pattern results from the presence of two dark-backed, chestnut-bellied subspecies (*D. brunneiventris vuilleumieri* and *D. b. brunneiventris*) that occur north and south, respectively, of the all-dark subspecies *D. humeralis aterrima* (Figure 1). Remsen (1984) proposed that leapfrog patterns are produced by random plumage mutations in the geographically intermediate taxon. Under this hypothesis, the all-black plumage of *D. humeralis atterima* would be due to a mutation that converts dark-back, chestnut-bellied birds into all-black birds. A testable prediction from this hypothesis is that the all-black mutant allele would appear as derived on a phylogeny of the *D. carbonaria* superspecies. Our SNAPP tree (Figure 2) suggests that *D. humeralis atterima* is nested within *D. b. brunneiventris*, and based on these results, the most parsimonious explanation of the all-black plumage in *D. humeralis atterima* is that it is a derived trait from a dark-backed, chestnut-bellied ancestor. Our data also suggest a more nuanced version of Remsen’s (1984) leapfrog hypothesis, including the underappreciated possibility that the movement of derived alleles via introgression could explain the distributional gap in leapfrog taxa.

### Implications for a more nuanced model of Andean biogeography

While the locations of phenotypic turnover between taxa (as indicated by dotted lines in Figure 1) correspond to known biogeographic regions (Graves 1982), which make sense in the context of habitat turnover and other north-south contact and suture zones (Cuervo 2013, Hazzi et al. 2018) in Andean birds, these did not always correspond to genetic turnover in the *D. carbonaria* superspecies. For example, the location of the genetic break we identified among individuals of *D. b. brunneiventris* did not overlap any of the other well-supported phylogeographic divides in the Andes (Hazzi et al. 2018). This genetic transition-zone does correspond to the range-limits of a few other high-Andean birds, such as the southern edge of the Rainbow Starfrontlet (*Coeligena iris*) range, the southern edge of the Many-striped Canastero (*Asthenes flammulata*) range, the northern edge of the Andean Negrito (*Lessonia oreas*) range, the northern edge of White-fronted Ground-Tyrant (*Muscisaxicola albifrons*) range, and the northern edge of the Glacier Finch (*Idiopsar speculifer*) range (Schulenberg et al. 2010), highlighting the lack of knowledge regarding biogeographic barriers to western-slope birds.

Moreover, our PCA results support the genetic distinctiveness of western-slope populations of *D. b. brunneiventris* in southwestern Peru, but they do not indicate any notable genetic break across the Marañon River Valley, the most notable biogeographic barrier for birds in the Andes (Chapman 1926, Parker et al. 1985, Winger et al. 2015), including other species of *Diglossa* (*D. cyanea*, Martínez-Gómez et al. 2023). Evidence of the genetic distinctiveness of taxa which have ranges that extend into northern Chile has been found in other avian groups (e.g., *Cranioleuca antisiensis*, Seeholzer and Brumfield 2018), which introduces questions regarding the commonality of this pattern. Perhaps these lineages should be described as distinct subspecies, or at a minimum designated as populations of conservation significance in terms of preserving the genetic diversity of Andean birds.

Future work in the region is warranted, but in the meantime, it seems clear that barriers (such as arid river valleys like the Marañon) for western slope birds and eastern slope humid cloud forest birds (Winger and Bates 2015, Winger 2017) are not necessarily the same. While latitude has long been identified as a correlate of isolation in montane taxa (Janzen 1967), disentangling the specific factors involved in these processes remains difficult because temperature and humidity co-vary with latitude. Identifying the barriers to high-elevation puna and scrub-associated birds of the western slope, as we did here, may provide insight into factors driving population divergence in montane regions. By comparing gene flow in taxa from the western and eastern slope of the Andes (or indeed, any mountain range with distinct assemblages on slopes), we can control for latitude, acting as a comparison point to examine the influence of differing humidity regimes due to the rain shadow effect (as occurs in the Andes) on the accumulation of the striking biodiversity found in mountainous regions across the globe (Myers et al. 2000).

## Supplementary material

Supplementary material is available at XXX online.

## Acknowledgements

We are indebted to the following museums and people who provided tissue loans for this work: the American Museum of Natural History (AMNH - Paul Sweet & Tom Trombone); University of Kansas Biodiversity Research Institute (KU - Mark Robbins); Louisiana State University Museum of Natural Science (LSU - Donna Dittmann, Van Remsen, & Fred Sheldon); Museum of Southwestern Biology (MSB - Christopher Witt & Andrew Johnson), Florida Museum of Natural History (UF - Andy Kratter), University of Alaska Museum (UAM - Kevin Winker). Thank you to Jessie Salter for assistance with lab work.

## Author contributions

A.E.H. conceived the idea for the study, undertook the analyses and data visualization, and wrote the manuscript. B.C.F conducted data analyses and curation. R.T.B and B.C.F edited the manuscript and supervised the research. All authors contributed to conceptualization, funding acquisition, and methodology.

## Data availability

The data and code that support this research are freely available at: XXX.

## SUPPLEMENT

**Table S1.** List of *Diglossa* individuals included in this study, the research collection in which the sample is archived, and the coordinates for the samples. Museum codes are as follows; AMNH = American Museum of Natural History, LSUMZ = Louisiana State University Museum of Natural Science, MSB = Museum of Southwestern Biology, KU = University of Kansas, UAM = University of Alaska Museum. The five individuals in bold were dropped from analyses during the data filtering stage.

| Species | Country | Museum ID | Lat | Long |
| --- | --- | --- | --- | --- |
| *albilatera* | Venezuela | AMNHDOT5024 | NA - outgroup | NA - outgroup |
| *albilatera* | Peru | LSUMZB44249 | NA - outgroup | NA - outgroup |
| *brunneiventris* | Peru | LSUMZB3578 | -9.77 | -76.06 |
| *brunneiventris* | Peru | LSUMZB72390 | -9.75 | -76.10 |
| *brunneiventris* | Peru | LSUMZB72539 | -7.78 | -77.75 |
| *brunneiventris* | Peru | LSUMZB72544 | -7.78 | -77.75 |
| *brunneiventris* | Peru | LSUMZB72550 | -7.78 | -77.75 |
| *brunneiventris* | Peru | LSUMZB7716 | -9.74 | -76.15 |
| *brunneiventris* | Peru | LSUMZB826 | -7.78 | -77.58 |
| *brunneiventris* | Peru | LSUMZB8299 | -9.93 | -75.77 |
| *brunneiventris* | Peru | LSUMZB8312 | -9.92 | -75.78 |
| *brunneiventris* | Peru | MSB34852 | -8.75 | -78.05 |
| *brunneiventris* | Peru | MSB34979 | -9.57 | -77.85 |
| *brunneiventris* | Peru | MSB36080 | -8.84 | -77.93 |
| *brunneiventris* | Peru | UF47413 | -6.87 | -78.09 |
| *brunneiventris* | Bolivia | AMNHDOT2758 | -15.13 | -68.89 |
| *brunneiventris* | Bolivia | AMNHDOT2773 | -15.13 | -68.89 |
| *brunneiventris* | Bolivia | AMNHDOT2893 | -14.83 | -68.94 |
| *brunneiventris* | Bolivia | AMNHSKIN834061 | -14.83 | -68.94 |
| ***brunneiventris*** | **Bolivia** | **AMNHSKIN834063** | **-14.83** | **-68.94** |
| *brunneiventris* | Peru | KU113967 | -13.33 | -74.20 |
| *brunneiventris* | Peru | KU115508 | -14.49 | -69.28 |
| *brunneiventris* | Peru | KU115509 | -14.49 | -69.28 |
| *brunneiventris* | Peru | KU131652 | -13.25 | -73.50 |
| *brunneiventris* | Peru | MSB27008 | -13.34 | -75.32 |
| *brunneiventris* | Peru | MSB27011 | -13.34 | -75.32 |
| *brunneiventris* | Peru | MSB27033 | -11.49 | -74.90 |
| *brunneiventris* | Peru | MSB27077 | -13.63 | -71.72 |
| *brunneiventris* | Peru | MSB27161 | -13.25 | -72.17 |
| *brunneiventris* | Peru | MSB31520 | -11.63 | -76.43 |
| *brunneiventris* | Peru | MSB31815 | -11.98 | -74.93 |
| *brunneiventris* | Peru | MSB31824 | -11.98 | -74.93 |
| *brunneiventris* | Peru | MSB33107 | -13.19 | -72.23 |
| *brunneiventris* | Peru | MSB33431 | -11.78 | -76.53 |
| *brunneiventris* | Peru | MSB33646 | -14.41 | -73.09 |
| *brunneiventris* | Peru | MSB34014 | -13.48 | -72.98 |
| *brunneiventris* | Peru | MSB34161 | -14.06 | -73.01 |
| *brunneiventris* | Peru | MSB35904 | -14.17 | -73.32 |
| *brunneiventris* | Peru | UAM25882 | -13.50 | -71.95 |
| *brunneiventris* | Peru | MSB35079 | -17.23 | -70.26 |
| *brunneiventris* | Peru | MSB35477 | -17.32 | -70.25 |
| *brunneiventris* | Peru | MSB35553 | -15.81 | -72.67 |
| *carbonaria* | Bolivia | LSUMZB106752 | -17.50 | -65.67 |
| *carbonaria* | Bolivia | LSUMZB1261 | -16.30 | -67.80 |
| ***carbonaria*** | **Bolivia** | **LSUMZB1275** | **-16.30** | **-67.80** |
| ***carbonaria*** | **Bolivia** | **LSUMZB1289** | **-16.30** | **-67.80** |
| *carbonaria* | Bolivia | LSUMZB1290 | -16.30 | -67.80 |
| ***carbonaria*** | **Bolivia** | **LSUMZB1294** | **-16.30** | **-67.80** |
| *carbonaria* | Bolivia | LSUMZB1295 | -16.30 | -67.80 |
| *carbonaria* | Bolivia | LSUMZB1300 | -16.30 | -67.80 |
| *carbonaria* | Bolivia | LSUMZB1306 | -16.30 | -67.80 |
| *carbonaria* | Bolivia | LSUMZB1309 | -16.30 | -67.80 |
| *carbonaria* | Bolivia | LSUMZB68138 | -17.47 | -65.87 |
| *carbonaria* | Bolivia | LSUMZB94721 | -17.84 | -64.73 |
| *carbonaria* | Bolivia | LSUMZB94723 | -17.84 | -64.73 |
| *carbonaria* | Bolivia | LSUMZB94728 | -17.84 | -64.73 |
| *carbonaria* | Bolivia | LSUMZB94736 | -17.84 | -64.73 |
| *carbonaria* | Bolivia | LSUMZB94751 | -17.84 | -64.73 |
| *carbonaria* | Bolivia | LSUMZB94752 | -17.84 | -64.73 |
| *carbonaria* | Bolivia | LSUMZB94753 | -17.84 | -64.73 |
| *humeralis* | Peru | LSUMZB31687 | -5.68 | -79.31 |
| ***humeralis*** | **Peru** | **LSUMZB31784** | **-5.69** | **-79.25** |
| *humeralis* | Peru | LSUMZB32036 | -5.69 | -79.25 |
| *humeralis* | Peru | LSUMZB32414 | -5.69 | -79.25 |
| *humeralis* | Peru | LSUMZB32730 | -5.69 | -79.25 |
| *humeralis* | Peru | LSUMZB415 | -5.36 | -79.55 |
| *humeralis* | Peru | LSUMZB419 | -5.36 | -79.55 |
| *humeralis* | Peru | LSUMZB429 | -5.36 | -79.55 |
| *humeralis* | Peru | MSB28082 | -6.24 | -79.25 |
| *humeralis* | Peru | MSB28100 | -6.24 | -79.24 |
| *humeralis* | Peru | MSB28215 | -6.24 | -79.23 |
| *humeralis* | Peru | MSB31331 | -6.24 | -79.25 |
| *humeralis* | Peru | MSB31333 | -6.24 | -79.23 |
| *humeralis* | Peru | MSB31335 | -6.24 | -79.25 |

**Figure S1.**
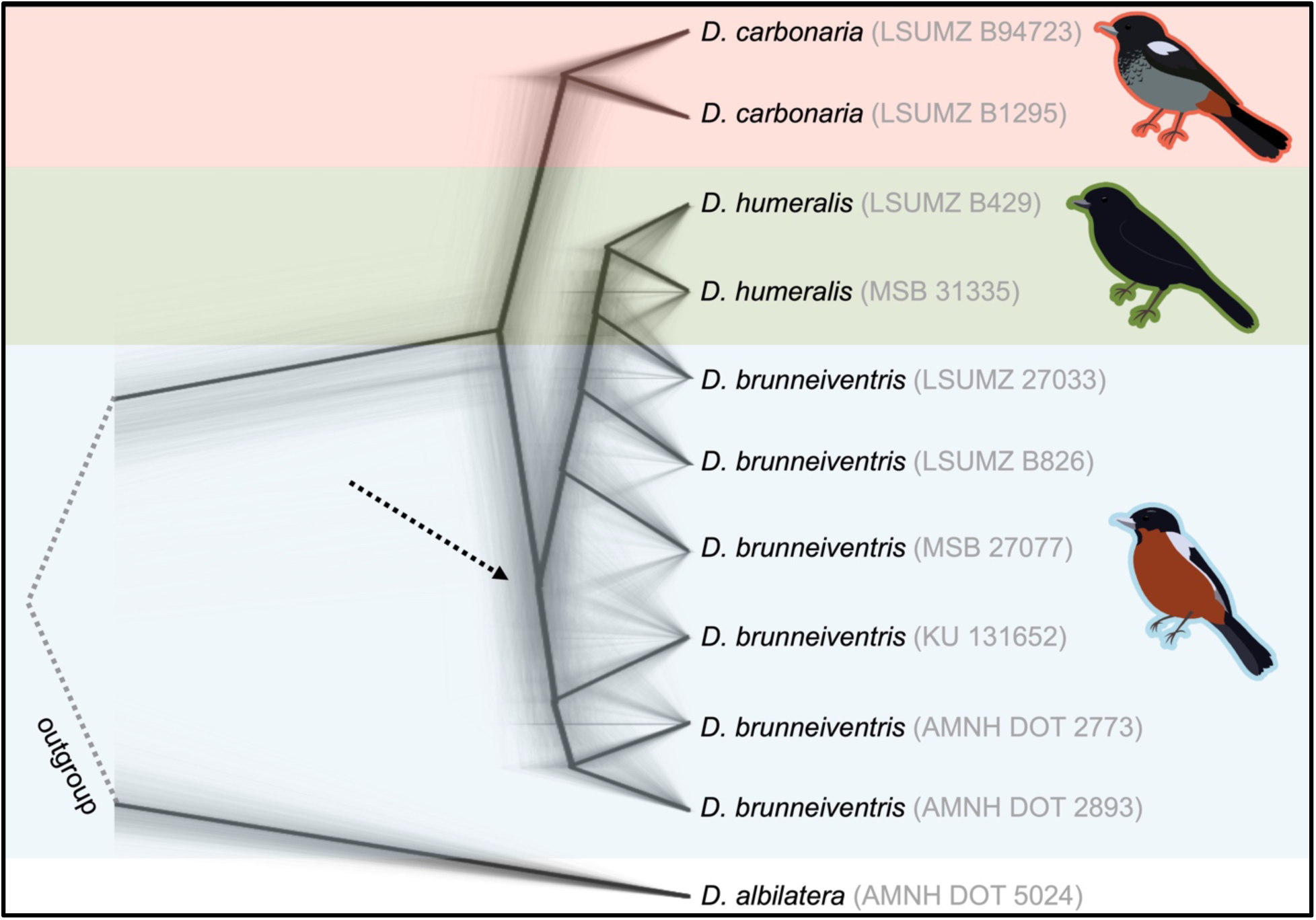
Densitree plots of a second SNAPP analysis showing the relationships of different individuals from each sampled population in the *D. carbonaria* superspecies, with *D. albilatera* as an outgroup (Figure 2 shows results from the first SNAPP analysis). The results are presented as a cloudogram of the posterior trees, with the density (or darkness) of the cloudogram indicating how often a particular topology occurs. Alternate topologies suggest uncertainty in the tree, particularly where indicated by the dotted arrows.

